# Panpipes: a pipeline for multiomic single-cell and spatial transcriptomic data analysis

**DOI:** 10.1101/2023.03.11.532085

**Authors:** Fabiola Curion, Charlotte Rich-Griffin, Devika Agarwal, Sarah Ouologuem, Tom Thomas, Fabian J. Theis, Calliope A. Dendrou

## Abstract

Single-cell multiomic analysis of the epigenome, transcriptome and proteome allows for comprehensive characterisation of the molecular circuitry that underpins cell identity and state. However, the holistic interpretation of such datasets presents a challenge given a paucity of approaches for systematic, joint evaluation of different modalities. Here, we present Panpipes, a set of computational workflows designed to automate multimodal single-cell and spatial transcriptomic analyses by incorporating widely-used Python-based tools to perform quality control, preprocessing, integration, clustering, and reference mapping at scale. Panpipes allows reliable and customisable analysis and evaluation of individual and integrated modalities, thereby empowering decision-making before downstream investigations.

## Background

Single-cell omics is a rapidly evolving field, with studies constantly expanding in size and scope to tackle increasingly complex biological and biomedical questions associated with development and ageing, health and disease, and vaccination and therapy [1–3]. Single-cell RNA sequencing (scRNA-seq) is now performed as a minimum, with a growing battery of methods becoming available to facilitate profiling of cell type-specific transcripts such as T- and B-cell receptor (TCR and BCR) repertoires (scV(D)J-seq) [4], as well as other molecular modalities, including chromatin states through single-cell sequencing Assay for Transposase-Accessible Chromatin (scATAC-seq) [5,6], and over 100 cell-surface protein markers using antibody-derived tags (ADT) for Cellular Indexing of Transcriptomes and Epitopes by sequencing (CITE-seq) [7].

Technological advances are also enabling experiments to move beyond the individual profiling of different molecular modalities and instead allow the simultaneous characterisation of the cellular genome, epigenome, transcriptome and/or proteome at a resolution that was previously inaccessible. To date, at least 25 different methods for the joint assaying of two or more modalities in single cells have been developed [3]. Single-cell multiomics are thus set to provide a fundamentally holistic understanding of cell and tissue circuitry and systems - surpassing insights that can be garnered from individual modalities alone.

Best practices for multimodal analysis are now emerging [8], with a wide range of packages and tutorials from which users can develop custom scripts [9,10]. For an end user, a typical analysis workflow could consist of collections of notebooks, which are run interactively and customised for each individual project. At different stages, the user is required to make choices about filtering strategies, normalisation, dimensionality reduction, and clustering, for example, in order to obtain a biologically meaningful interpretation of their data. However, this scenario does not constitute an efficient application of best practices: relying only on custom scripts poses a risk due to a lack of methodological consistency, thus jeopardizing reproducibility [11]. This problem is particularly relevant for large-scale projects, where sequential analysis rounds are necessary, as the dataset increases in size. Therefore, fully harnessing the power of multiomic single-cell technologies is impeded by the absence of comprehensive pipelines which seamlessly integrate best practices by jointly analysing modalities in a reproducible, automated and computationally efficient fashion.

To meet this need, we have developed Panpipes, a set of automated, flexible and parallelised computational workflows for the processing of multimodal single-cell data. Panpipes is implemented in Python, and has at its core the scverse foundational tools for single-cell omics data analysis [12]. It leverages the efficient and flexible data handling structures AnnData [13] and MuData [14], complemented by a number of widely-used single-cell analysis tools including Scanpy [15], muon [14], scvi-tools [16] and scirpy [17]. These packages have been successfully applied in a variety of settings, including the building of large-scale atlases and deep learning computational tasks, for instance as part of the Human Cell Atlas reference building efforts [18], and they scale to millions of cells, whilst maintaining reasonable computation times.

Single-cell analysis frameworks, such as Seurat [10] or Scanpy [15], have promoted the democratisation of access to single-cell data analysis. Seurat leverages R’s statistical capabilities, whilst Scanpy relies on Python’s machine learning libraries, and both use distinct data structures (SeuratObject and AnnData and MuData, respectively). Each framework has its own inherent strengths and they cater to different programming communities (largely R versus Python users). Interactively analysing single-cell data within a single framework can be useful for exploratory investigations and analysis of smaller datasets, but can pose challenges when testing multiple parameter combinations, especially for complex and large datasets. To meet these challenges, pipelines for single-cell analysis are emerging, which utilise Workflow Management Languages to orchestrate analysis with one or more frameworks. Such pipelines are designed for creating data processing workflows that automate and expedite complex processes involving multiple tasks and dependencies. Utilisation of pipelines can thus enable the parallel comparison of different algorithms and tools. This is critical as although benchmarking studies provide important guidance for algorithm or tool selection [19–24], no single method will necessarily generate the best results for all datasets, and benchmarking studies can also reach different conclusions [19,20].

Published pipelines for single-cell analyses such as scFlow [25], scrnaseq [26], bollito [27] and pipeComp [28] are restricted to single modality (RNA only) datasets, and typically use R-based packages such as Seurat [10] and Scater [29]. Other published pipelines are designed to run using cloud computing and employ web-based interfaces such as SCiAp [30], Granatum [31] or ASAP [32]. However, these web-based workflows can be restrictive in terms of analysis parameters and users may struggle with larger datasets. In contrast, Panpipes is designed to run on high-performance computing (HPC) clusters, but retains the capacity to be deployed locally for small datasets, providing the user with added run flexibility. The pipeline is managed using the Computational Genomics Analysis Tools (CGAT)-core framework [33], which simplifies and parallelises job submission both on local computers or by interacting with common cluster workflow managers such as SLURM [34].

Panpipes is the first set of open-source workflows for the analysis of multimodal single-cell and spatial transcriptomic datasets. Panpipes performs quality control, preprocessing, integration, clustering, reference mapping and spatial transcriptomics deconvolution at scale. The user’s interaction with Panpipes is highly customisable, enabling analysts to have fine control over their analyses, in a reproducible manner. Our pipeline is written in a modular way such that the workflows can be further developed in order to keep up with the fast-moving field of single-cell multiomics and spatial transcriptomics. As Panpipes leverages scverse tools which are interoperable between Python and R ecosystems, our choice of relying on scverse, which is a well-maintained community project, will ensure that Panpipes is future-proof.

## Results

### Panpipes: a pipeline for single-cell multiomic and spatial transcriptomic analysis

Panpipes comprises six workflows for the analysis of single-cell multiomic datasets: “Ingestion”, “Preprocessing”, “Integration”, “Clustering”, “Reference Mapping” and “Visualisation” (Fig. 1). Panpipes also includes four workflows dedicated to spatial transcriptomics, including: “Ingestion”, “Preprocessing”, “Clustering” and “Deconvolution” (Fig. 1). The unifying aim across these workflows is to guide the user through the key decision-making steps of the analytical process and to gather all the data necessary to annotate cell types and states.

**Figure 1.**
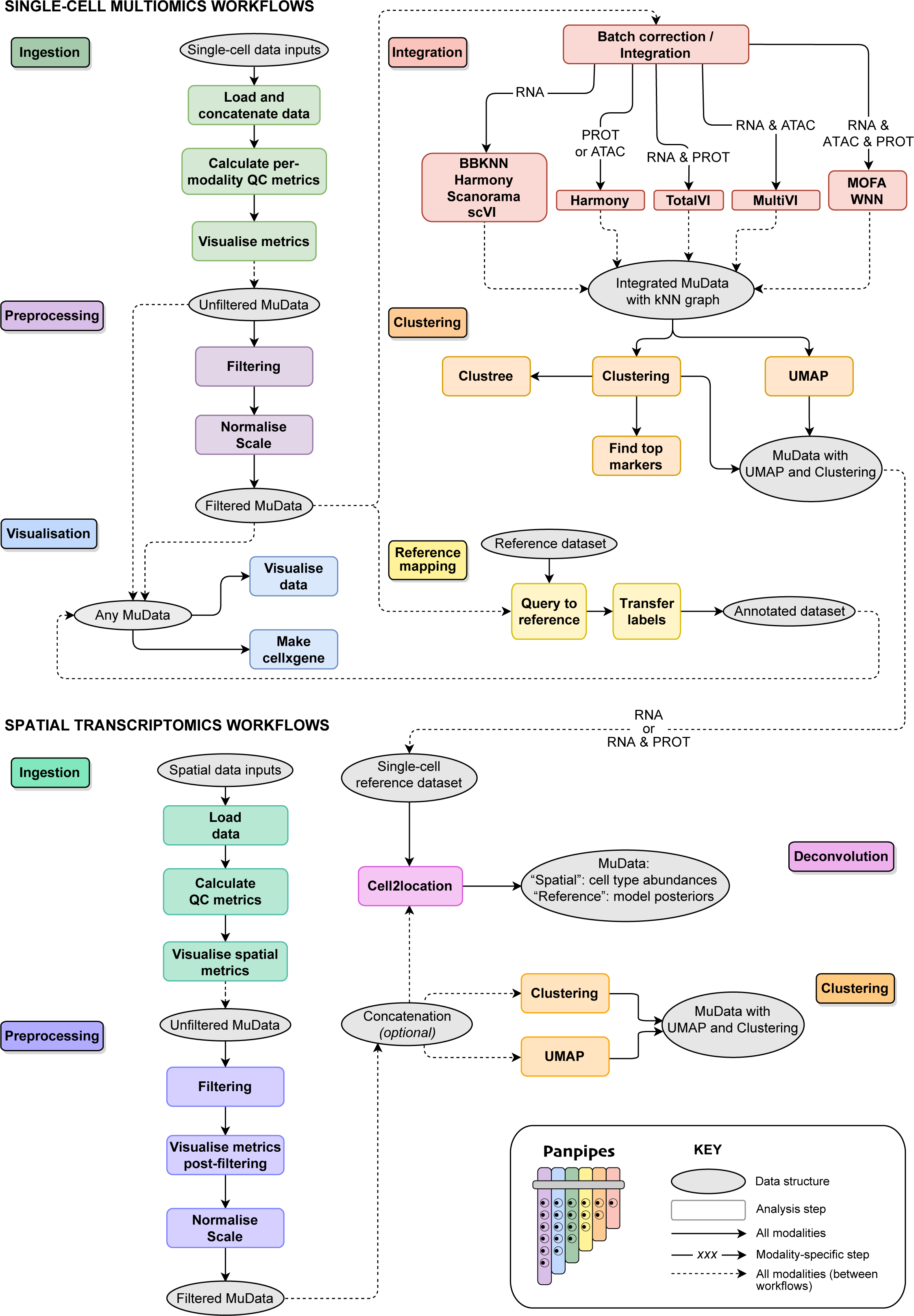
The Panpipes workflows for single-cell multiomic and spatial transcriptomic analysis. Panpipes has a modular design and performs ingestion, preprocessing, integration and batch correction, clustering, reference mapping and spatial transcriptomics deconvolution with custom visualisation of outputs. The schematic demonstrates the flow of data within (solid arrows) and between (dashed arrows) workflows, and modality-specific steps are indicated.

The single-cell multiomics “Ingestion” workflow leverages Scanpy and muon functionalities, together with custom functions, to ingest single-cell data from any combination of assays including gene expression, immune repertoire profiling, CITE-seq, and ATAC-seq. A variety of file formats can be used as input, namely count matrices, CellRanger outputs and h5 files. This flexibility simplifies the process of quickly analysing published datasets as well as novel data from any single-cell platform. Finally, the data are concatenated and saved as a MuData (h5mu) file, which is a multimodal container [14]. Additionally, the workflows can incorporate metadata associated either with the sample, such as patient-level information or with the barcode, such as demultiplexing information or cell-level annotations in the case of preprocessed data. Standard and custom QC metrics are computed and visualised for each modality.

The second stage of Panpipes, the "Preprocessing" workflow for the single-cell multiomic data is used to (i) filter the data based on previously computed quality control (QC) metrics, (ii) (optionally) downsample, (iii) normalise and (iv) scale the data, with different options available for each modality. After preprocessing, any of the following workflows can be run, depending on the analytical requirements: “Integration”, “Clustering”, and “Reference Mapping”. “Integration” is used to integrate and (optionally) batch correct via a choice of uni- and multimodal methods, which can be run in a parallel fashion. “Clustering” runs parallelised clustering over a wide range of parameters using dimension reductions from either “Preprocessing” or “Integration” as inputs. “Reference Mapping” utilises query-to-reference (Q2R) and label transfer (LT) functionalities from scvi-tools and single-cell architectural surgery (scArches) [35] to integrate and annotate query data with reference data.

Finally, “Visualisation” is included as a separate overarching workflow as the outputs from any of the other workflows can be used as its inputs. It produces a range of plots, combining the experiment-specific metadata and the analysis outputs from the other workflows, to aid the inspection and interpretation of results. Users who have run multiple methods and parameter choices in parallel can evaluate their results at each step and can export the final objects to cellxgene [36] for user-friendly and interactive exploration.

For spatial transcriptomics analyses, Panpipes’ “Ingestion” workflow leverages Scanpy and squidpy [37] functionalities to read data generated through the 10x Genomics’ Visium or Vizgen’s MERSCOPE platforms. After “Ingestion” the “Preprocessing” workflow is used to (i) filter the data, (ii) visualise and evaluate QC metrics post-filtering, and (iii) normalise and (iv) scale the data. Subsequently the data can be (optionally) concatenated and then “Clustering” is performed. For the 10x Genomics’ Visium data, whose resolution is dictated by the number of cells found over ‘spots’ containing spatially barcoded capture probes, a “Deconvolution” workflow can also be run after “Preprocessing”, which enables leveraging of single-cell references to computationally achieve a higher resolution of cell type identification within spots.

### Evaluation of single-cell multiomic data quality with Panpipes

To enable data QC and thus the identification and obtainment of high-quality cells, Panpipes generates a battery of metrics standard to the evaluation of scRNA-seq data [38,39], such as the total number of unique molecular identifiers (UMI) per cell-barcode and the percentage of UMIs assigned to mitochondrial transcripts. In addition, users can provide custom gene lists to score specific cellular phenotypes. This can facilitate the retention of cell types with more atypical properties such as plasma cells or neutrophils [40], that might otherwise be excluded. It also renders Panpipes compatible with any genome, thereby enabling analyses of cells from other species.

In addition to RNA-associated metrics, Panpipes produces a number of QC visualisations which are specific to ATAC-seq assays (ATAC) or ADT assays (PROT), or are related to the joint QC of multiple modalities (Fig. 2). For ATAC, the fragment and barcode metrics are incorporated in the data object and the nucleosome signal is computed. With the inclusion of a peak annotation file which maps chromosome coordinates to gene IDs, transcription start site enrichment is also calculated. For PROT, comparing the UMI counts in the cell-containing foreground against the empty droplets in the background can give an indication of whether antibodies are binding specifically, or contributing to ambient contamination in the dataset (Fig. 2A). The level of the background staining in empty droplets on a per ADT basis in the PROT assay, correlates with the signal strength of the ADTs after normalisation, and thus is likely to influence downstream analysis. Panpipes provides two PROT normalisation options, centred log ratio transformation (CLR) [7] and denoised and scaled by background (dsb) normalisation [41]. CLR generates a natural log ratio of the count for a protein in a cell relative to other cells, hence enabling improved distinction of cell populations, but without endeavouring to account for background or technical noise [7]. The dsb normalisation aims to correct for ambient ADTs and unspecific binding of antibodies to cells [41]. Panpipes allows for the normalised PROT expression profile to be visually inspected for individual ADTs via histograms for each normalisation method (Fig. 2B, C), whilst scatter plots facilitate head-to-head comparisons of the methods on a per ADT basis (Fig. 2D). In addition, Panpipes QC can profile the ambient fractions of RNA and PROT expression data, to provide insight into the variation of the background relative to the foreground across samples for both modalities (Fig. 2E).

**Figure 2.**
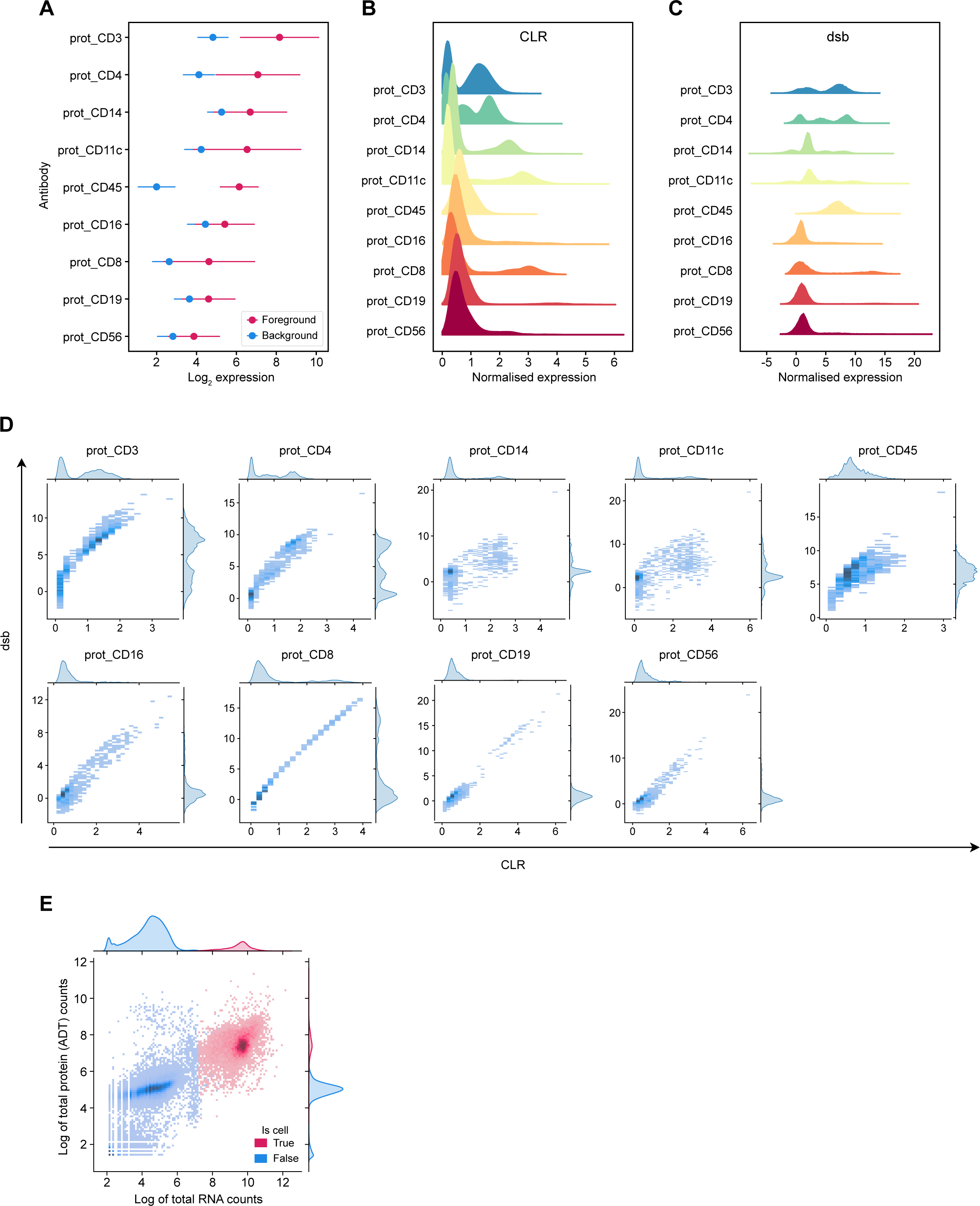
Cell-surface protein QC metric visualisations generated by Panpipes. **A** Expression (log_2_ of raw counts) of cell-surface protein markers in cells (foreground, red) versus empty droplets (background, blue). Dots represent mean expression and lines show the standard error of the mean. **B** Distribution of cell-surface protein marker expression after CLR normalisation. **C** Distribution of cell-surface protein marker expression after dsb normalisation. **D** Joint distribution plot of cell-surface protein marker expression normalised by CLR (x-axis) versus dsb (y-axis). **E** Joint distribution plot of RNA (x-axis) versus the cell-surface protein (encoded by the RNA; y-axis) in cells (red) versus empty droplets (blue). Expression of the RNA and protein is plotted as the log of the total counts (per cell barcode).

The capacity to extensively inspect QC metrics for all modalities present in a single-cell dataset, is critical for subsequent clustering, annotation and downstream analyses [42], and can help inform decision-making with respect to multimodal integration.

### Multimodal integration for unified cellular representation

Following QC, Panpipes offers a parallelised framework to aid the user in choosing a reduced dimensionality representation of a given dataset based on a unimodal or multimodal integration, with the option to apply batch correction to individual modalities or in a joint fashion.

To mimic a typical analysis scenario in which a user may wish to apply different processing choices simultaneously, we demonstrate Panpipes’ functionality on a trimodal dataset (TEA-seq) [43] of three samples with joint single-cell measurements of RNA, PROT and ATAC. The workflow enables each individual modality to be projected onto a latent representation with or without a selection of batch correction methods [44,45] (e.g. BBKNN for RNA, Harmony for PROT, and BBKNN and Harmony for ATAC as shown in Fig. S1). The batch correction methods offered for the different modalities have been selected based on underlying statistical assumptions and published benchmarks [20]. Multimodal batch-aware integration methods can also be employed for two or more modalities, including MultiVI (used for RNA+ATAC with this dataset) [46], totalVI (for RNA+PROT) [47], and weighted nearest neighbour (WNN; for ATAC+RNA+PROT) [10] (Fig. 3A-E). MultiVI and totalVI perform multimodal integration whilst accounting for batch covariates while WNN affords the highest processing flexibility as it can perform multimodal integration after individual modalities are individually batch corrected. Users are provided with a choice of unimodal and multimodal integration tools as each integration approach may answer a different biological question, depending on the dataset. The variation in the performance of these tools for batch merging can be visualised through UMAP representations (Fig. 3A-E) and is also evaluated by the calculation of Local Inverse Simpson’s Index (LISI) scores (Fig. 3F) [45].

**Figure 3.**
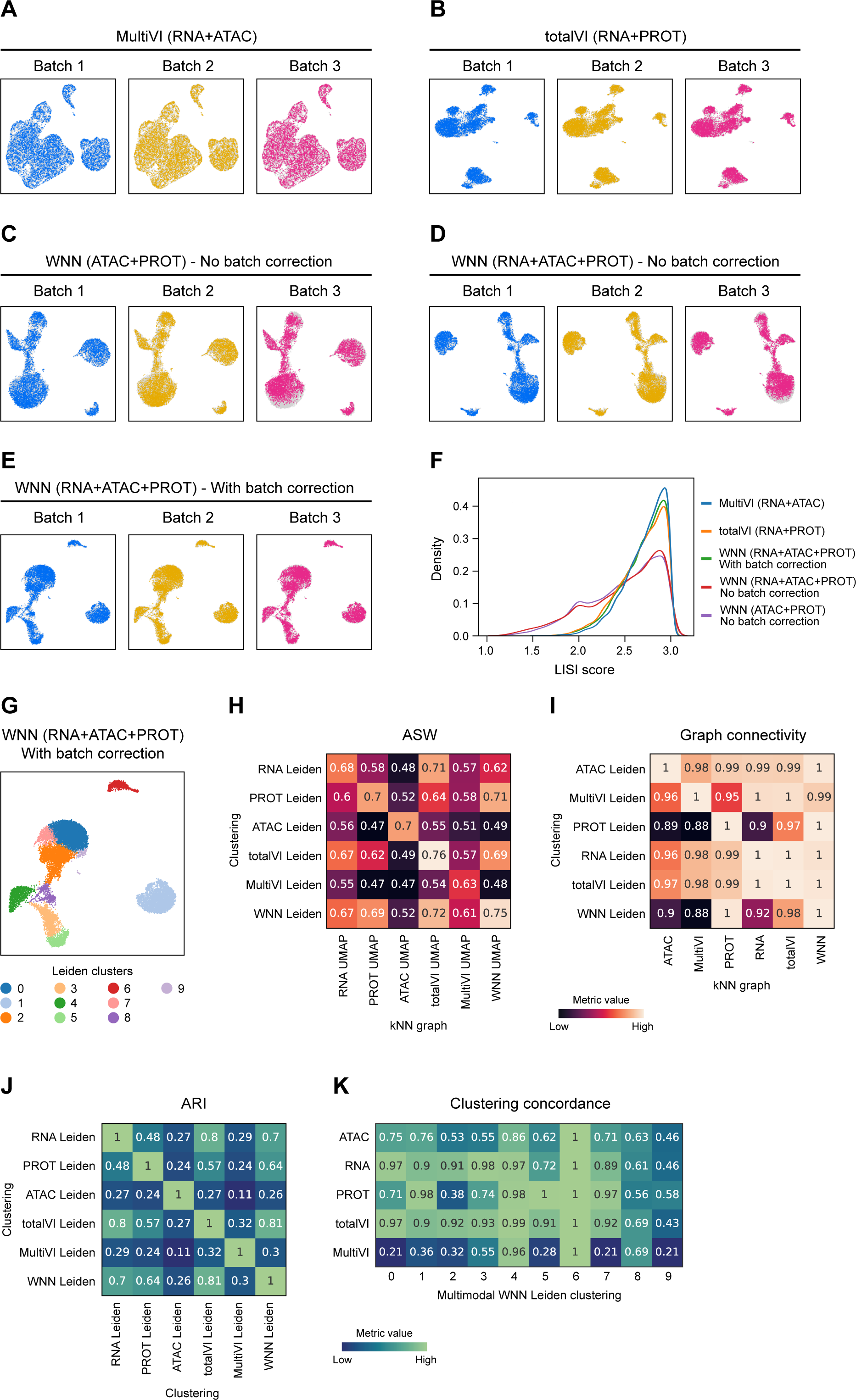
Panpipes integration workflow enables evaluation of multimodal integration and batch correction. UMAPs showing individual batches (batch 1, blue; batch 2, ochre; batch 3, pink) after RNA and ATAC modality integration using MultiVI (**A**), RNA and ADT cell-surface protein (PROT) integration using totalVI (**B**), ATAC and PROT integration using WNN with no batch correction (**C**), RNA, ATAC and PROT integration with no batch correction (**D**) and with batch correction (**E**). **F** LISI score distribution for integrations depicted in **A-E**. **G** UMAP of Leiden clustering after trimodal WNN integration with batch correction. **H** ASW metric values for different clustering labels and cell embeddings. **I** Graph connectivity metric values for different clustering labels and cell embeddings. **J** ARI metric values for cluster matching with different clustering choices. **K** Per-cluster concordance of clustering choices relative to trimodal WNN Leiden clustering.

After integration, each unimodal or multimodal embedding is clustered using the “Clustering” workflow (Fig. 3G) and further integration evaluation is carried out using a selection of single-cell integration benchmarking (scIB) metrics [20] and custom visualisations (Fig. 3H-K). Average silhouette width (ASW) and graph connectivity [48] are applied on an integrated object with a choice of clustering labels and cell embeddings (Fig. 3H, I). These metrics estimate how well similar cells cluster together by considering intra- and inter-cluster similarities and local connectivities, respectively, with higher scores signifying better performance. Since each clustering returns a cell partitioning from the embedding it was generated on, it may be anticipated that any single tested clustering would have the highest score for its original embedding; however, this is not always the case. For example, with the TEA-seq dataset, comparing the RNA clustering with the totalVI embedding and the PROT clustering with the WNN embedding yielded similarly high or higher ASW and graph connectivity scores (Fig. 3H, I). As specific multimodal integration metrics have not been developed yet, this demonstrates how Panpipes’ repurposed use of scIB metrics in the multimodal scenario is instrumental in identifying where individual modalities may have uneven contributions to the final cell classification.

To further assess the concordance of clustering choices calculated from different modalities, Panpipes generates a cluster matching metric, the Adjusted Rand Index (ARI) [49], for global concordance evaluation (Fig. 3J). Panpipes also implements another clustering concordance visualisation on a per-cluster basis, whereby one clustering choice is selected as the reference (in the example, multimodal WNN clustering; Fig. 3K). For each of the clusters identified in the reference, the extent to which alternative approaches provide at least one cluster that groups together the same cells as the reference, is then scored. Higher scores indicate that a high percentage of the cells in the reference cluster are also grouped together in the alternative cell partitioning. With the TEA-seq dataset for instance, WNN cluster 6 is entirely recapitulated by all the alternative clustering choices, whilst cluster 9 is poorly represented by the alternatives (all scores <0.60; Fig. 3K).

Thus, Panpipes provides the user with the capacity to efficiently run and thoroughly evaluate the correction of batch effects and the integration of individual and multiple modalities to facilitate the selection of the optimal integration method prior to downstream analyses.

### Reference mapping with Panpipes

Leveraging the “Reference Mapping” workflow, Panpipes also has reference mapping capabilities. As large-scale single-cell multiomic datasets become increasingly available [1,50], users will wish to take advantage of such resources to expedite cell annotation of their own data and aid biological interpretation. However, learning from reference datasets can pose an analytical challenge due to batch effects, computational resource limitations and data access restrictions [26]. Panpipes can aid building unimodal or multimodal references and enables the user to query multiple references simultaneously using scArches [16,35,51]. For example, a user can perform filtering of low-quality cells on the input dataset (via “QC” and “Preprocessing”) and can then immediately run the “Reference mapping” workflow without proceeding with the “Integration” and “Clustering” workflows. Alternatively, users can annotate their query dataset independently, then project it onto a reference and evaluate concordance with the reference labels. The concordance of the transferred labels with the original labels is evaluated in the query via a selection of scIB metrics. Furthermore, users can leverage Panpipes to query the same dataset on multiple references, allowing for comparison between them.

To demonstrate the “Reference mapping” workflow we have performed Q2R and LT using as the query a unimodal scRNA-seq peripheral blood mononuclear cell (PBMC) dataset [52] and three PBMC references varying in size and in the granularity of cell type labels. These references include one RNA-specific unimodal dataset (PBMC_R1) [53] and two multimodal PBMC datasets (PBMC_R2 and PBMC_R3) [10]. Single-cell Annotation using Variational Inference (scANVI) [54] and totalVI were employed for the uni- and multimodal references, respectively (Fig. 4).

**Figure 4.**
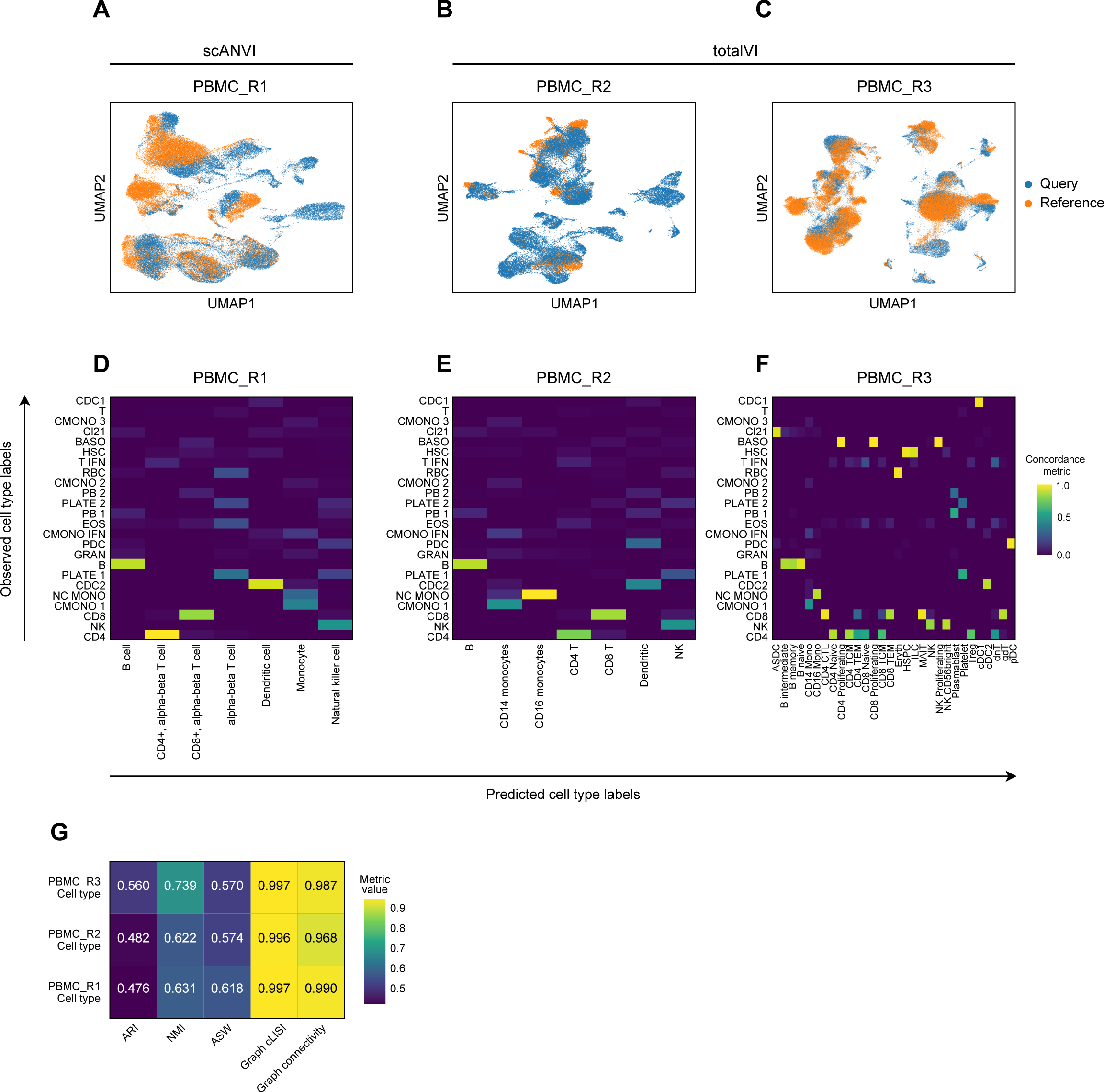
Panpipes reference mapping allows evaluation of multiple reference datasets. **A** UMAP of unimodal Q2R integration between the query dataset (orange) and the PBMC_R1 reference dataset (blue) using scANVI. **B** UMAP of multimodal Q2R integration between the query dataset (orange) and the PBMC_R2 reference dataset (blue) using totalVI. **C** UMAP of multimodal Q2R integration between the query dataset (orange) and the PBMC_R3 reference dataset (blue) using totalVI. **D** Query to PBMC_R1 label transfer concordance with predicted (reference-derived) labels on the x-axis and observed (query-derived) labels on the y-axis. **E** Query to PBMC_R2 label transfer concordance with predicted (reference-derived) labels on the x-axis and observed (query-derived) labels on the y-axis. **F** Query to PBMC_R3 label transfer concordance with predicted (reference-derived) labels on the x-axis and observed (query-derived) labels on the y-axis. **G** Label conservation scoring using scIB metrics for each Q2R integration.

Q2R integration is visually assessed by UMAP plots (Fig. 4A-C), whilst LT performance is evaluated by estimating the concordance of predicted and observed labels (Fig. 4D-F). Notably, for the datasets tested, there was variation in the cells present in the query relative to the reference data, and this was detectable by both the incomplete integrations by inspecting the UMAP generated on query and reference latent embeddings, and by the imperfect LT concordance (Fig. 4A-F). Interestingly, with reference dataset PBMC_R3 [10], a query cluster annotated as basophils (“BASO”) received three different proliferating lymphoid cell reference labels suggesting that the outputs generated by Panpipes can help to identify annotation inconsistencies for further investigation and thus obtain an optimal annotation. Finally, label conservation is scored using metrics that assess local neighbourhoods, (including graph cLISI and graph connectivity), global cluster matching, (including ARI and normalised mutual information (NMI) [55]), and relative distances as determined by cell-type ASW (Fig. 4G).

The capacity of Panpipes to employ and compare multiple reference datasets will be critical as single-cell omics atlases continue to expand in scale and complexity and users will likely want to draw upon all resources available to arrive at a high-confidence annotation of their own data.

### Orchestrating spatial transcriptomic analysis

The rapid evolution of spatial transcriptomics technologies allows us to capture gene expression within the context of tissue architecture [56–59]. Similar to the Panpipes single-cell workflows, the spatial transcriptomics workflows also include “Ingestion”, “Preprocessing” and “Clustering”, and enable the parallel analysis of data derived from multiple spatial transcriptomics slides.

Critically for the 10x Genomics Visium ‘spot’-based approach, whereby the data for each individual RNA capture area (‘spot’) will represent a mixture of transcriptomes from all the cells found in the area, a “Deconvolution” workflow is provided. This is based on the use of the cell2location Bayesian model that integrates single-cell and spatial transcriptomic data to effectively resolve the transcriptomes from each capture area into finer cell types [60]. The “Deconvolution” workflow can utilise external single-cell datasets, but also seamlessly integrates with the single-cell multiomics workflows to utilise single-cell data generated subsequent to the Panpipes single-cell “Integration” and “Clustering” (Fig. 1).

### Benchmarking

To demonstrate Panpipes’ performance, we ran the “Integration” workflow on six datasets of different sizes, representing the full data and subsamples of a PBMC dataset [10] and the TAURUS study gut dataset [61], assessing runtime (Fig. 5A) and resource usage (Fig. 5B,C). Since Panpipes implements each integration method as an independent component, the main advantage of our pipeline is the management of data flow and the possibility to choose which method to run in a parallel fashion, allowing the independent processing of modalities across multiple methods (Fig. 5).

**Figure 5.**
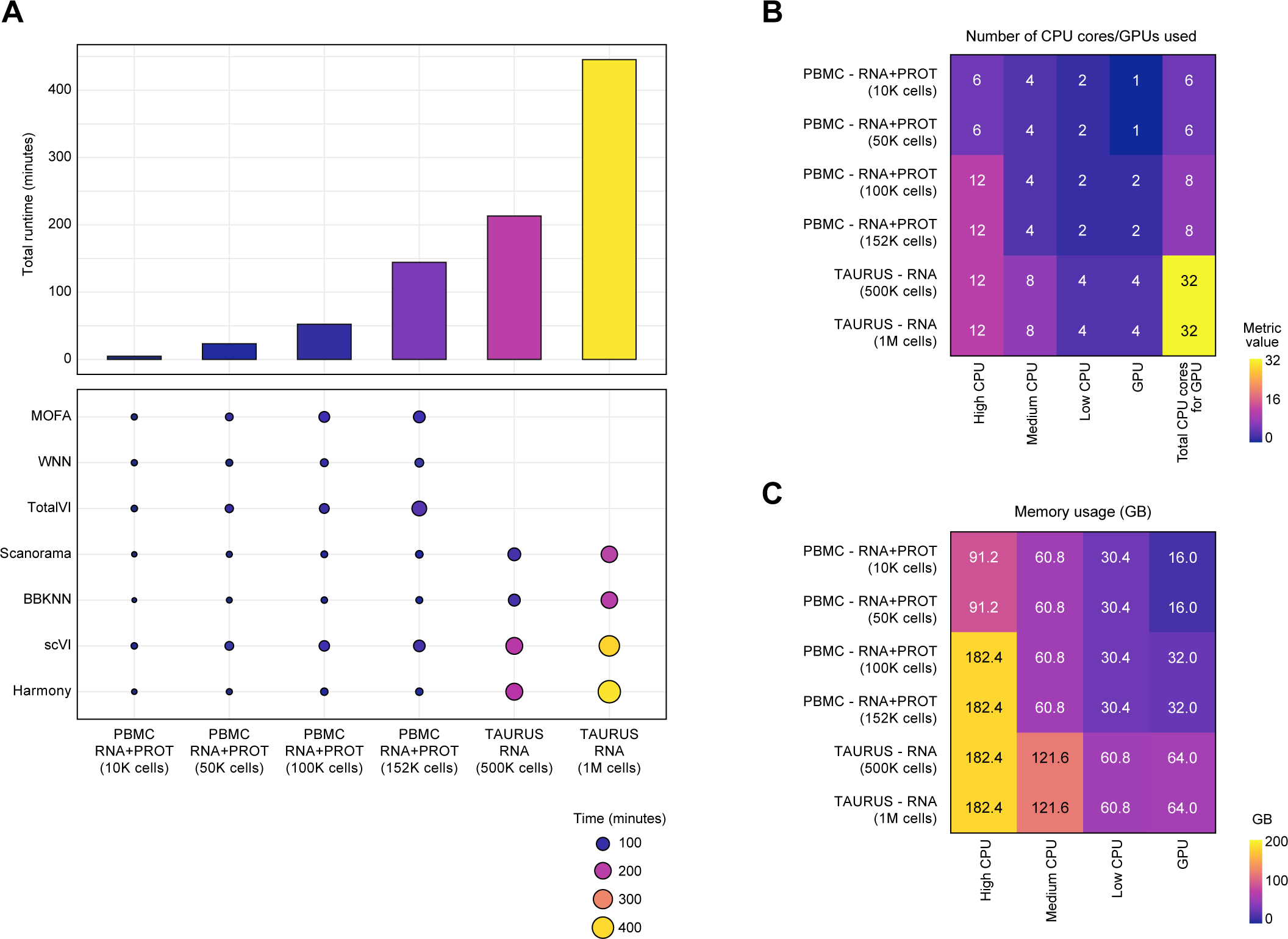
Panpipes time and resource usage benchmarking. **A** Total runtimes (bar plot) and run times by integration method (dot plot) when using the “Integration” workflow on six different datasets representing the full data and subsamples of a PBMC CITE-seq dataset and the TAURUS study gut scRNA-seq dataset. **B** Resource usage as denoted by the number of central processing unit (CPU) cores or graphics processing units (GPUs) utilised for the six datasets. **C** Memory usage as denoted by the number of GB utilised for the six datasets.

Furthermore, Panpipes’ outputs provide a biological perspective for evaluation and interpretation of the data in any biological context. For example, taking the PBMC data [10] used for the time and resource benchmarking, for which cell type annotations are available, we ran the “Integration” workflow in a multimodal, integration batch-aware fashion, with the aim of assessing which of the multimodal integration methods included in Panpipes was able to recapitulate the ground truth annotation. Assessment of integrations generated by MOFA, totalVI and WNN by the UMAP plot distribution of the cell type labels (Fig. S2) shows, a good separation of the cell types with every method, although with MOFA two batches of each cell type could be observed suggesting poor integration. However, with totalVI and WNN these batches were not discernible, but WNN (with harmony correction of RNA and PROT) resulted in the clearest separation of the CD4+ and CD8+ T and natural killer (NK) cell subsets, for example (Fig. S2).

## Discussion

We have developed Panpipes, a highly flexible pipeline to perform finely-tuned analyses on both single modality and multiomic single-cell datasets. Panpipes is based on scverse [12], which includes the most popular Python-based single-cell packages, and thus readily scales up to millions of cells. Panpipes allows the ingestion, quality checking, filtering, integration, clustering, reference mapping and visualisation of complex single-cell datasets. Our workflow can process any single-cell dataset containing RNA, cell-surface proteins, ATAC and immune repertoire modalities, as well as spatial transcriptomics data generated through the 10x Genomics’ Visium or Vizgen’s MERSCOPE platforms.

Panpipes is designed to help analysts run a comprehensive evaluation of their single-cell data. The pipeline does not stop at basic preprocessing but instead constitutes a flexible framework to explore more complex analytical choices including unimodal and multimodal integration with or without batch correction. Panpipes implements nine different integration methods, helping the user select the best parameterization for their specific analytical needs. The reference mapping functionality offered by Panpipes can expedite cell annotation and provides a powerful setting to evaluate the utility of multiple reference datasets. This may be particularly important given that individual references may not perfectly recapitulate the biological or biomedical context of the query dataset. Moreover, given the agility of Panpipes, the pipeline may be well suited to benchmarking studies, in particular in relation to multimodal integration and reference mapping, which are currently lacking in the field of single-cell multiomics.

We have developed Panpipes with a wide range of users in mind. The pipeline is publicly available with extensive documentation and tutorials which allow users to efficiently go from raw data to clustering and annotation in a semi-automated fashion – regardless of whether they are using unimodal, multimodal or spatial transcriptomic datasets. Being Python-based, Panpipes could represent an easy entry point for users with a Machine Learning background who have more limited single-cell analysis expertise. Our pipeline may also appeal to core computational facilities in academia or in the pharmaceutical industry that need a quick and flexible single-cell pipeline that readily allows for the assessment of common problems such as ambient molecular contamination and batch effects, and that facilitates the utilisation of external single-cell datasets to help inform target identification and evaluation.

Panpipes is also FAIR principle [62] compliant, in line with the requirements of many funding agencies. The source code is easily finable and accessible on GitHub (https://github.com/DendrouLab/panpipes) and as a PyPi package, and further documentation and tutorials (https://panpipes-pipelines.readthedocs.io/en/latest/) are provided to encourage users to adapt the pipeline to their own needs.

## Conclusions

The last few years have seen a continuous and rapid development of multimodal protocols that scale to millions of cells and thousands of analytes in the single-cell omics field. The collection of analytical methods that deal with the complexity of large single-cell datasets is likely to increase, with a marked interest in methods that allow integration of multiomic assays [63].

Given this fast evolution of the single-cell and also the spatial omics analysis landscape, Panpipes is in continuous development. Panpipes is modular by design to enable its extension to incorporate new methods that can deal with further omics modalities in the future. These could include single-cell genomic DNA sequencing and epigenome profiling beyond chromatin accessibility, and other technologies such as flow cytometry, mass spectrometry and hyperplexed imaging. Panpipes provides a platform for both customisation and reproducibility of single-cell multiomic and spatial transcriptomic analyses, ensuring a stable foundation for the consistency and continuity of scientific discovery.

## Methods

### Implementation Details

Panpipes comprises workflows implemented using the CGAT-core framework [33]. CGAT-core automates submission to and parallelisation of jobs across HPC clusters. Flexible environment control is implemented using conda. To interact with the pipeline, the user is required to simply edit a YAML file for each workflow to customise the parameters for their own analyses. Finer details of these options are listed below. We provide documentation on each workflow and how to run them in https://panpipes-pipelines.readthedocs.io.

### Ingestion

Data from various sources is ingested to be combined and formatted as a MuData object. Specific QC metrics are computed for each modality, following guidelines defined in single-cell best practices [8]. Scrublet is used to compute doublet scores [64]. Cells are also scored based on custom gene lists (e.g. mitochondrial and ribosomal gene proportions). Gene lists compatible with human and mouse are provided and users can readily input features for alternative species or define their own QC metrics based on custom gene lists.

### Preprocessing

The thresholds determined by the QC pipeline outputs are included as parameters in the YAML file, and the data are filtered accordingly. In the “Preprocessing” workflow, the user is able to specify custom filtering options on any set of metrics computed in the QC workflow. Next, for the RNA data, the data are normalised and scaled, and the highly variable genes are computed using Scanpy functionalities. In parallel, PROT data are normalised using either CLR [7] or dsb [41] using muon functionalities and functions implemented ad hoc. For example, users can specify which margin to normalize the PROT data to, namely by cell or within the features’ distribution. ATAC data are log normalised or normalised by term frequency-inverse document frequency, following the options offered by the muon package.

### Integration

“Integration” implements a range of algorithms in order to batch correct individual modalities, and to combine multiple modalities in a low-dimensional space. For each unimodal processing, the dimensionality reduction of choice (PCA and/or Latent Sematic Indexing (LSI) for ATAC) is applied and the data are batch corrected based on user-defined parameters. Four unimodal batch correction algorithms are included in Panpipes: BBKNN [44], Harmony [45], Scanorama [65], and scVI [16,51]. Panpipes supports both modality-specific multi-modal integration batch-aware methods such as MultiVI for ATAC and gene expression, and totalVI [47] for PROT and gene expression, and modality-agnostic methods such as MOFA [66] and WNN [10]. The results of these integrations are compared with the aid of scIB metrics [20], inspection of LISI scores [45] and visual inspection of UMAP plots.

### Clustering

“Clustering” implements both Leiden and Louvain clustering of a connectivity graph constructed on a reduced dimension computed in the “Integration” workflow. The reduced dimension data can be a single modality representation e.g. PCA or Harmony components, or a multi-modality representation e.g. MultiVI or totalVI reduced dimension. The clusters are then visualised on a UMAP computed from the same dimensionality reduction, or the user has the option to project clusters onto any of the computed UMAPs from alternative dimensionality reductions. The user can compute clustering for a wide range of resolutions, to quickly assess the cell type representation within their dataset. Cluster assignments across different resolutions are compared using clustree [67]. Finally, the workflow calculates the top multimodal markers for each computed clustering, offering a choice of different statistical tests for the scoring of the features based on Scanpy’s rank_genes_groups().

### Reference mapping

The “Reference mapping” workflow implements Q2R and LT from scvi-tools and scArches supported models, namely scVI, scANVI and totalVI models. Code is implemented with the scvi-tools package. Data for query and reference datasets can be supplied as individual AnnData [13] or MuData [14] objects, and reference models generated with any of the aforementioned methods. The user is required to specify a minimal set of mandatory parameters and can specify additional covariates and define custom training parameters by customising the pipeline.yml.

### Visualisation

The “Visualisation” workflow is implemented to aid inspection and interpretation of results. The visualisation workflow uses matplotlib, seaborn and ggplot to generate boxplots, histograms, scatterplots, and dimensionality reduction plots (such as PCA or UMAPs), using any combination of variables across the modalities, and experimental metadata. The “Visualisation” workflow is also used to export the data objects to cellxgene [36] for interactive visualisation. Importantly, this cellxgene object contains UMAP plots from multiple modalities so that the user can directly review gene, protein, peak expression and repertoire information on the same set of UMAPs.

### Ingestion (spatial)

The “Ingestion_spatial” workflow is implemented to ingest data from various spatial transcriptomics platforms such as 10x Genomics’ Visium or Vizgen’s MERSCOPE. Multiple slides can be processed in parallel. Similar to the single-cell “Ingestion” workflow, the spatial data are quality controlled following best practices recommendations. This workflow produces a MuData object with a “spatial” layer and the newly generated QC values.

### Preprocessing and Clustering (spatial)

Similar to the single-cell workflow, “Preprocessing” for the spatial data follows the “Ingestion” workflow to allow filtering and processing of the spatial data. Custom QC parameter thresholds are included in the YAML file, and the data are filtered accordingly. Next, the data are normalised and scaled, and the highly variable genes are computed using Scanpy functionalities. Finally, dimensionality reduction is run and saved to the MuData “spatial” object. The output of the spatial “Preprocessing” workflow can be run through the spatial “Clustering” which is as described for the single-cell multiomics workflow but with additional parameters for spatial transcriptomic data.

### Deconvolution

The “Deconvolution” workflow allows the inference of cell type composition of ‘spot’-based spatial transcriptomic data, using a single-cell reference. “Deconvolution” implements cell2location [60] and can be run on multiple individual slides with the same single-cell reference.

### Processing of data for figures

Uni- and multimodal processing of the trimodal TEA-seq data

Data for the trimodal TEA-seq dataset was obtained from [43]. Briefly, the three raw datasets for each individual modality were each concatenated into a unimodal AnnData object. ATAC fragment indexes were regenerated using Tabix [52], and the peaks of the three batches were merged following the signac tutorial [53]. The three objects were then partitioned to the cell barcodes in common across the modalities and fed to the “QC” pipeline as individual AnnData objects, which produced a unified MuData container for the three modalities. QC and filtering were performed independently on each modality and the intersection of the passing QC cells across the three modalities was chosen for further analysis. Cells were integrated using unimodal or multimodal integration methods as described (Fig. 4 and Fig. S1).

### Reference mapping

We mapped the same query PBMC dataset [52] to three different PBMC references [10,53]. For the scANVI example, we produced the reference model using scVI and then updated the model to scANVI to leverage the label transfer functionality. The query was then mapped to the reference data using the “Reference mapping” workflow by specifying the batch covariate and the cell type label. For the TotalVI example, we used the data presented in https://docs.scvi-tools.org/en/stable/tutorials/notebooks/totalVI_reference_mapping.html. The second reference dataset (PBMC_R2) was obtained by downloading the 10x Genomics PBMC 5k and 10k datasets presented in https://scarches.readthedocs.io/en/latest/totalvi_surgery_pipeline.html.

The reference was generated using the “Integration” workflow calling the totalVI algorithm, and cells were labelled using the expression of the protein surface markers. We generated the third reference dataset (PBMC_R3) following the process described in the scvi-tools tutorial, downloading the data using the internal scVI function adata_reference = scvi.data.pbmc_seurat_v4_cite_seq(mask_protein_batches=5). To enable transferring the labels to the query from both PBMC_R2 and PBMC_R3, a Random Forest classifier was trained on the latent TotalVI embedding of the reference model. All query and reference datasets were provided to the “Reference mapping” workflow to perform Q2R and LT, specifying batch covariates and cell type labels, training the query model with number of epochs = 200 and leaving all other default parameters.

### Benchmarking

A PBMC CITE-seq dataset [10] and the TAURUS study gut scRNA-seq dataset [61] were used for benchmarking. For these analyses, both the full datasets and the downsampled datasets (10K, 50K, and 100K cells for the PBMCs and 500K cells for the gut cells) were utilised. The code used for the benchmarking is available at https://github.com/DendrouLab/panpipes-benchmarks.

## Availability of data and materials

### Code Availability

Source code, full documentation and tutorials are available at https://github.com/DendrouLab/panpipes and https://panpipes-pipelines.readthedocs.io. Panpipes maintenance and updates are the responsibility of the co-senior authors Calliope A. Dendrou and Fabian J. Theis and co-first author Fabiola Curion.

### Single-cell Datasets

The data used in Fig. 2 to showcase the ADT-associated metrics were obtained from https://www.10xgenomics.com/resources/datasets/10-k-human-pbm-cs-with-total-seq-b-human-tbnk-antibody-cocktail-3-v-3-1-3-1-standard-6-0-0. The trimodal TEA-seq dataset was downloaded using dbGAP accession number phs002316 [43]. For reference mapping analyses, the PBMC CITE-seq dataset used as the query was obtained from the Gene Expression Omnibus (GEO) under accession number GSE155673 [52]. The PBMC datasets used as references are available as follows: CITE-seq data from [10] available via GEO (accession number GSE164378) and dbGAP (accession number phs002315.v1.p1), and scRNA-seq data from [53] available via covid19cellatlas.org. The PBMC data (accession number GSE164378) were also used in Fig. 5 and the gut data for this figure were obtained from the TAURUS study [61].

## Acknowledgments

The authors thank Dr Melissa Grant-Peters, Mr Wojciech Lason, and Dr Jacqueline Siu for testing the code and identifying bugs. The authors also wish to thank Dr Luke Zappia for revising the manuscript and for helpful discussions. This work has been performed within the framework of the Cartography Consortium (https://www.medsci.ox.ac.uk/for-staff/resources/business-partnerships-office/oxford-janssen-working-in-collaboration), which authors CAD, CRG and DA are members of, and which is funded by Janssen Biotech, Inc.

## Author information

### Contributions

FC and CRG conceived the study with input from CAD and FJT. FC and CRG wrote the code with contributions from TT and DA and SO. FC and CRG wrote the manuscript with input from CAD and FJT. FC, CRG, CAD and FJT read and revised the manuscript. FC and CRG are equal contributors to this work, and can reference this work as a first authorship paper in their curriculum vitae.

## Ethics declarations

### Ethics approval and consent to participate

Not relevant to our study.

### Ethics consent for publication

Not relevant to our study.

## Supplementary figure legend

**Figure S1. Panpipes integration workflow enables evaluation of batch correction of different individual modalities.**

UMAPs showing individual batches (batch 1, blue; batch 2, ochre; batch 3, pink) with no correction or after unimodal Harmony or BBKNN batch correction for individual (**A**) RNA, (**B**) ADT cell-surface protein (PROT), and (**C**) ATAC modalities from the trimodal TEA-seq dataset. LISI score distribution for no batch correction (blue) or unimodal Harmony (green) or BBKNN (orange) batch correction for individual (**D**) RNA, (**E**) PROT, and (**F**) ATAC modalities from the trimodal TEA-seq dataset.

**Figure S2. Panpipes integration workflow enables visualisation and evaluation of integration and batch correction in a biological context.**

UMAPs showing the integration of a previously annotated, available PBMC CITE-seq dataset after utilisation of MOFA, totalVI, and WNN. The distribution of the PBMC subsets in the UMAP plots can be visually inspected, taking into consideration the cell type labels. DCs: dendritic cells; NK: natural killer.

