## Supplementary figures and images for "Panpipes: a pipeline for multiomic single-cell and spatial transcriptomic data analysis"

### Supplementary Figure S1

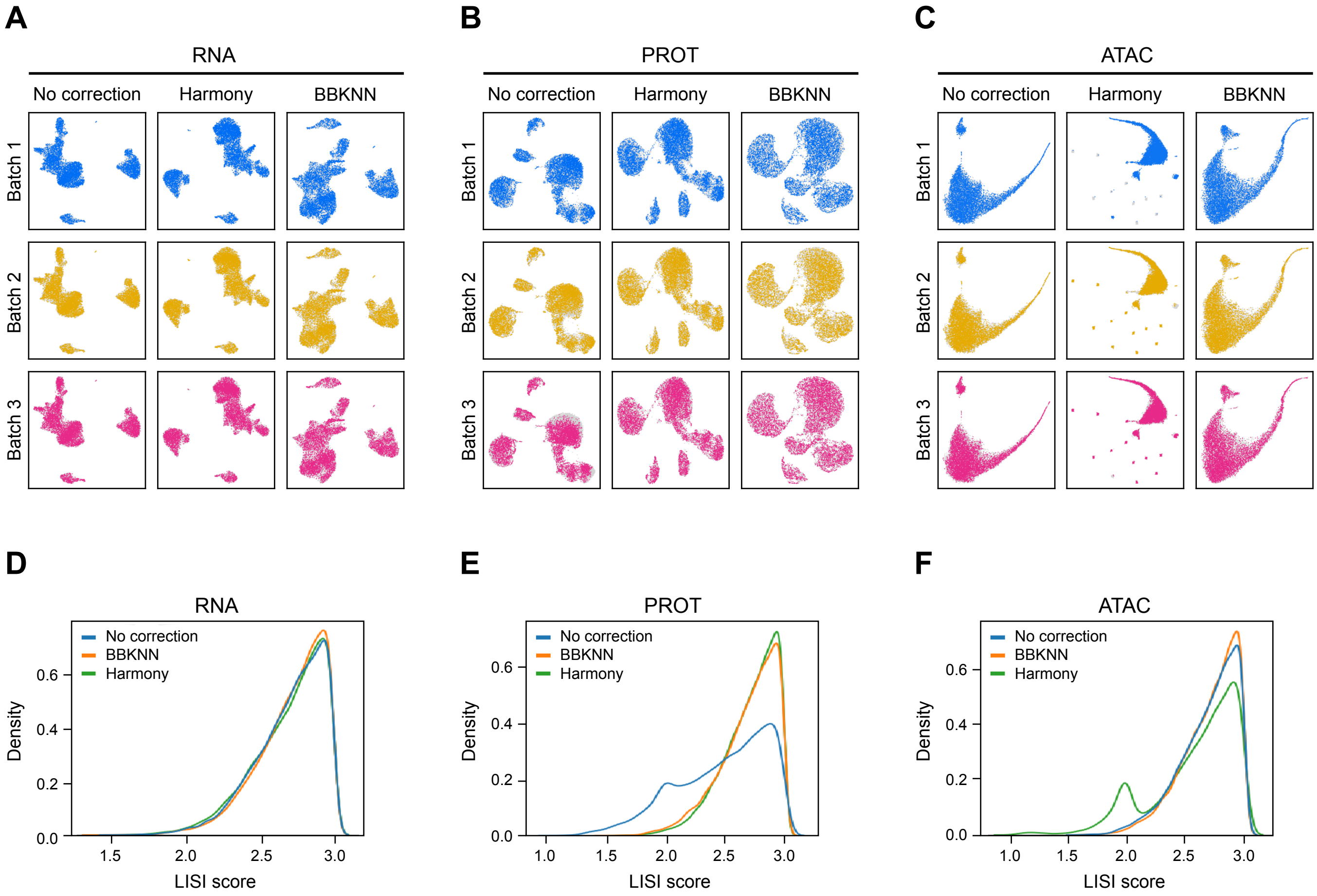

### Supplementary Figure S2

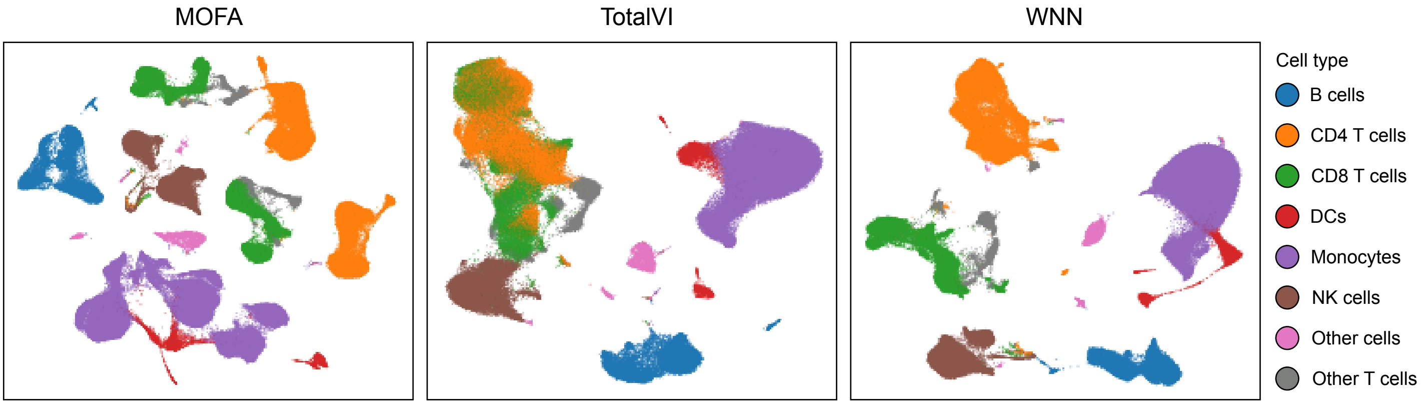
